# Functional module detection through integration of single-cell RNA sequencing data with protein–protein interaction networks

**DOI:** 10.1101/698647

**Authors:** Florian Klimm, Enrique M. Toledo, Thomas Monfeuga, Fang Zhang, Charlotte M. Deane, Gesine Reinert

## Abstract

Recent advances in single-cell RNA sequencing (scRNA-seq) have allowed researchers to explore transcriptional function at a cellular level. In this study, we present scPPIN, a method for integrating single-cell RNA sequencing data with protein–protein interaction networks (PPINs) that detects active modules in cells of different transcriptional states. We achieve this by clustering RNA-sequencing data, identifying differentially expressed genes, constructing node-weighted PPINs, and finding the maximum-weight connected subgraphs with an exact Steiner-tree approach. As a case study, we investigate RNA-sequencing data from human liver spheroids but the techniques described here are applicable to other organisms and tissues. scPPIN allows us to expand the output of differential expressed genes analysis with information from protein interactions. We find that different transcriptional states have different subnetworks of the PPIN significantly enriched which represent biological pathways. In these pathways, scPPIN also identifies proteins that are not differentially expressed but have a crucial biological function (e.g., as receptors) and therefore reveals biology beyond a standard differentially expressed gene analysis.

## 1 Introduction

Liver metabolism is at the centre of many non-communicable diseases, such as diabetes and cardiovascular disease [1]. In healthy organisms, the liver is critical for metabolic and immune functions and gene-expression studies have revealed a diverse population of distinct cell types, which include hepatocytes in diverse functional cell states [2]. As diabetes is a complex and heterogenous disease, the study of liver physiology at single-cell resolution helps us to understand the biology [3]. At a single-cell level, however, large-scale protein interaction data is not yet available [4]. In this study, we develop scPPIN a method for the integration of single-cell RNA-sequencing data with complementary PPINs. Our scPPIN analysis of liver single cell data and PPINs reveals biological pathways in cells of different transcriptional states that hint at inflammatory processes in a subset of hepatocytes.

In recent years, much attention has been given to scRNA-seq techniques as they allow researchers to study and characterise tissues at a single-cell resolution [5, 6, 7]. Most importantly, scRNA-seq reveals that there exist clusters of cells with similar gene expression profiles, commonly referred to as ‘cell states’ [8]. Multiple approaches have been created to reveal these cell clusters, driven by the transcriptional profile of each cell [9, 10]. The analysis of differentially expressed genes (DEGs) between these cell clusters has been shown to reveal different cell types [11], diseased cells [12], and cells that resist drug treatment [13]. Due to technological advances the quality and availability of scRNA-seq data has increased dramatically in the last decade [14]. This makes the development of computational approaches for interpreting scRNA-seq data an active field of research [15] of which one research direction is the identification of gene regulatory networks in scRNA-seq data (e.g., SCENIC [16], PIDC [17]).

These approaches do not make systematic use of available protein–protein interaction data. One can represented such data as PPINs and use PPINs to, for example, identify essential proteins [18, 19, 20] and to predict disease associations [21, 22] or biological functions [23, 24, 25]. For this, researchers have used tools from network science and machine learning. Many of these methods build on the well-established evidence that in PPINs, proteins with similar biological functions are closely interacting with each other. These groups of proteins with common biological functions are called *modules* [26, 27].

It is understood that gene-expression is context-specific and thus varies between tissues [28], changes over time [29], and differs between healthy and diseased states [30]. It follows therefore that different parts of a PPIN are active under different conditions [31]. Analysing PPINs in an integrated way, together with bulk gene-expression data, provides such biological context, helps to reveal context-specific active functional modules [32, 33], and can identify proteins associated with disease [34].

Based on the success of methods where PPINs have been integrated with bulk expression data, we have developed scPPIN, a novel method to integrate scRNA-seq data with PPINs. It is designed to detect active modules in cells of different transcriptional states. We achieve this by clustering scRNA-seq data, performing a DEGs analysis, constructing node-weighted PPINs, and identifying maximum-weight connected subgraphs with an exact Steiner-tree approach. Our method is applicable to the broad range of organisms for which PPINs are available [35].

The scPPIN method can be used to analyse mRNA-seq data from any tissue or organ type. As a case study, we investigate scRNA-seq data from human liver spheroids because this tissue is important in many diseases and it is known to have diverse cell types with different cellular metabolic processes. This makes the application of our method particularly relevant, because we expect the identification of very different active modules in different cell clusters — a hypothesis that our investigation partially confirms.

Our method identifies proteins involved in liver metabolism that could not be detected from the scRNA-seq data alone. Some of them have been shown to be important in the liver of other organisms and for others this study is the first indicator of important function in liver. Furthermore, we can associate cells in a given transcriptional state with enriched biological pathways. In particular, we find that cell clusters have different biological functions, for example, translational initiation, defence response, and extracellular structure organisation.

This case study demonstrates that scPPIN provides insights into the context-specific biological function of PPINs. Importantly, these insights would not have been revealed from either data type (PPIN or scRNA-seq) alone. We therefore anticipate that this technique will reveal novel insights for other organisms and tissue types.

## 2 Results

In this paper, we present scPPIN, a method that detects functional modules in different cell clusters. The method involves multiple analysis steps (see Fig. 1 for an overview and the Method Section for a detailed discussion). First, we preprocess the scRNA-seq profiles. Second, we use an unsupervised clustering technique from Seurat to identify sets of cells in similar transcriptional states. Third, for each cluster, we identify DEGs using a Wilcoxon rank sum test. Fourth, for each gene in every cluster, we compute additive scores from these p-values (see [32] and Supplementary Note 1). Fifth, for each cluster, we map these gene scores to their corresponding proteins in a PPIN, constructed from publicly available data from BioGRID [35]. Lastly, we identify functional modules as maximum-weight connected subgraphs in these node-weighted networks.

**Figure 1:**
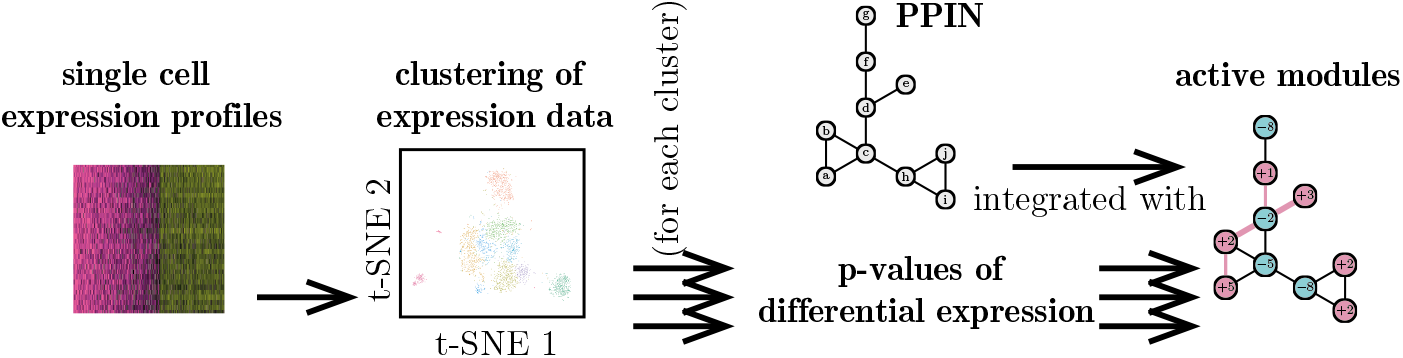
Our method consists of the following steps. (1) clustering of scRNA-seq data (e.g., with Seurat [9]). For each cluster, we (2) compute p-values of differential expression and use them to (3) estimate node scores by using an approach presented in [32]. (4) We combine these node scores with a PPIN to construct node-weighted PPINs for each cluster. (5) We compute functional modules as maximum-weight connected subgraphs.

In order to demonstrate scPPIN, we investigate newly measured scRNA-seq data of liver hepatocytes (see Method Section 8 for a description of the experimental setup and preprocessing steps). Using a standard modularity-maximisation algorithm, we obtain ten cell clusters of which six consist of hepatocytes (see Fig. 2), which make up a majority of the liver tissue. Hepatocytes are known to show a functional diversity and are, for example, important for the carbohydrate metabolism [2]. We focus on hepatocytes because it allows us to study the heterogeneity of cellular function in this single cell type.

**Figure 2:**
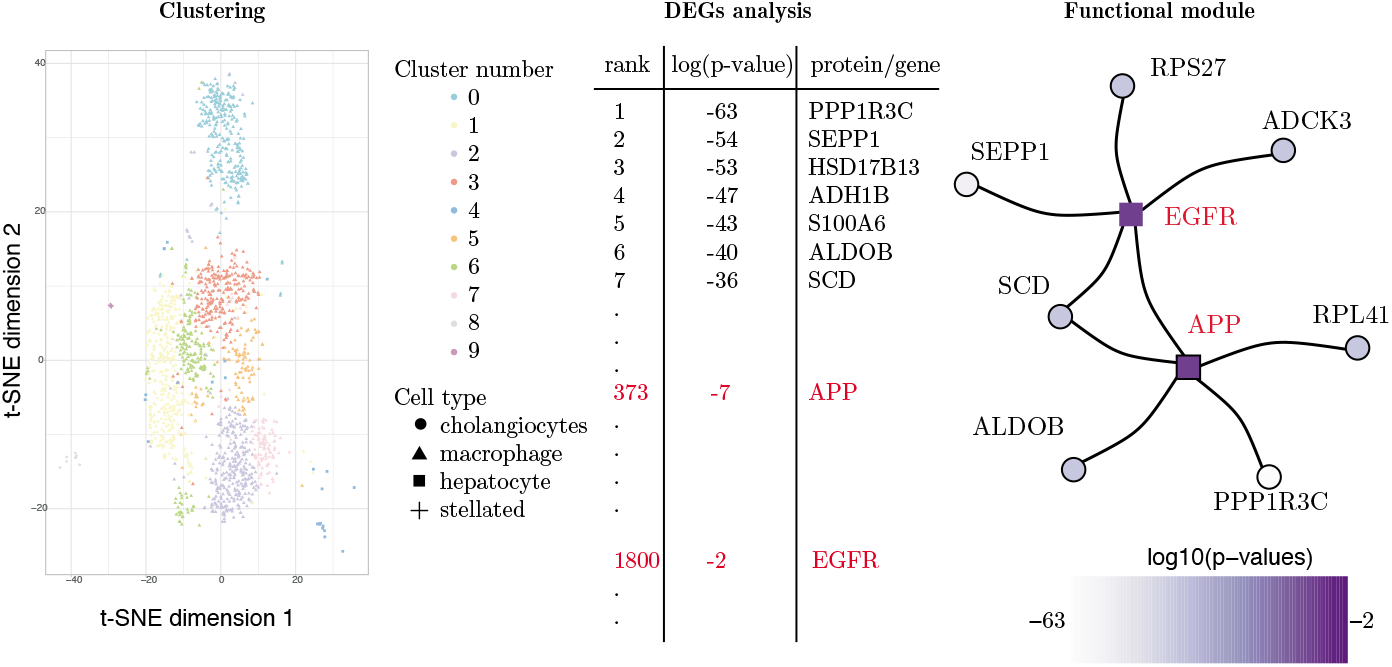
(Left) Clustering the scRNA-seq data reveals ten clusters of which six are hepatocyte cells. We visualise the cells in a two-dimensional space as obtained from a *t-distributed stochastic neighbour embedding* (t-SNE) of their original high-dimensional space [36]. (Middle) For each cluster, we perform DEGs analysis to identify genes that most differentially expressed in a given cluster. Here, we show DEGs for hepatocyte cluster 6 (H6). (Right) We use scPPIN to integrate the p-values from DEGs analysis with the PPIN and identify a functional module for the H6 cluster. We find genes that are significantly differentially expressed (disks) and proteins that are not strongly differentially expressed (squares). Colour indicates p-value from low (white) to high (purple).

We identify DEGs in each of the hepatocyte clusters. In Fig. 2, we show the p-values of differential expression for some of the genes in hepatocyte cluster H6. Usually, the top-ranked genes in each of the clusters can be seen as ‘marker genes’, i.e., one may use these genes to associate cells with a certain transcriptional state. For H6, for example, *protein phosphatase 1 regulatory subunit 3C* (PPP1R3C) has with 10^-63^ the smallest of all p-values. It therefore could serve as a potential biomarker and is a known regulator of liver glycogen metabolism [37]. While such a DEG analysis reveals important genes in certain cell states it is not straightforward to identify the crucial biological pathways. Next, we demonstrate that integrating p-values from a DEGs analysis with PPIN information can reveal a more comprehensive picture of the biological processes. In Fig. 2, we show a functional module identified by scPPIN. We detect a subnetwork consisting of nine proteins. This module consist of seven proteins with small p-values (among them PPP1R3C) that are connected to each other via the *amyloid precursor protein* (APP) and *epidermal growth factor receptor* (EGFR), which have p-values ~ 10^-7^ and 10^-2^, respectively. Both proteins are integral membrane proteins and do not show significant differential expression in this cell cluster as they rank 373 and 1800 out of all differentially expressed genes. The EGFR signalling network has been identified as a key player in liver disease [38]. The precise function of APP is unknown but it is involved in Alzheimer’s disease and also has been hypothesised to be involved in liver metabolism [39].

These findings demonstrate that scPPIN can help to automate the further investigation of results from a DEGs analysis by identifying parts of the PPIN that correspond to genes that are significantly differentially expressed. Furthermore, it also identifies proteins corresponding to genes that are not significantly differentially expressed in a particular cluster. These genes are candidates of a biological connector function between differentially expressed genes.

### 2.1 Influence of the False Discovery Rate

We have demonstrated that scPPIN can reveal functional modules inside a PPIN and associate them with cells of a certain transcriptional state. Now we explore whether there is only one functional module for a given cell state or rather functional modules of different sizes.

There is only one free parameter in scPPIN, the false discovery rate (FDR). Intuitively, increasing the FDR identifies a larger subgraph of the PPIN as an active module. In the following, we explore this systematically, for the hepatocyte cluster H6 that we investigated above.

The size *M* ∈ [1,*N*] of the detected modules is non-decreasing with the FDR. While the size *M* is non-decreasing, our method is non-monotonous, i.e., proteins identified for a certain FDR are not necessarily detected for all larger FDRs. For small FDRs, we detect a module of size *M* =1, which is exactly the protein with the smallest p-value^1^. For FDRs close to one, we detect a maximum weight subgraph which is spanning almost the whole network.

In Fig. 3, we show the size M of the the optimal subnetworks for cluster H6 as a function of the FDR. As expected, the *M*(FDR) is non-decreasing. For FDR < 10^-26^, we detect a single node, which represents PPP1R3C, the protein with the smallest p-value (~ 10^-63^). For larger FDRs, we detect subnetworks of larger size that contain proteins that are associated with larger p-values and could not have been identified with the gene-expression data alone. For FDR = 10^-25^, for example, we detect the subnetwork of size *M* = 9 (shown in Fig. 2).

**Figure 3:**
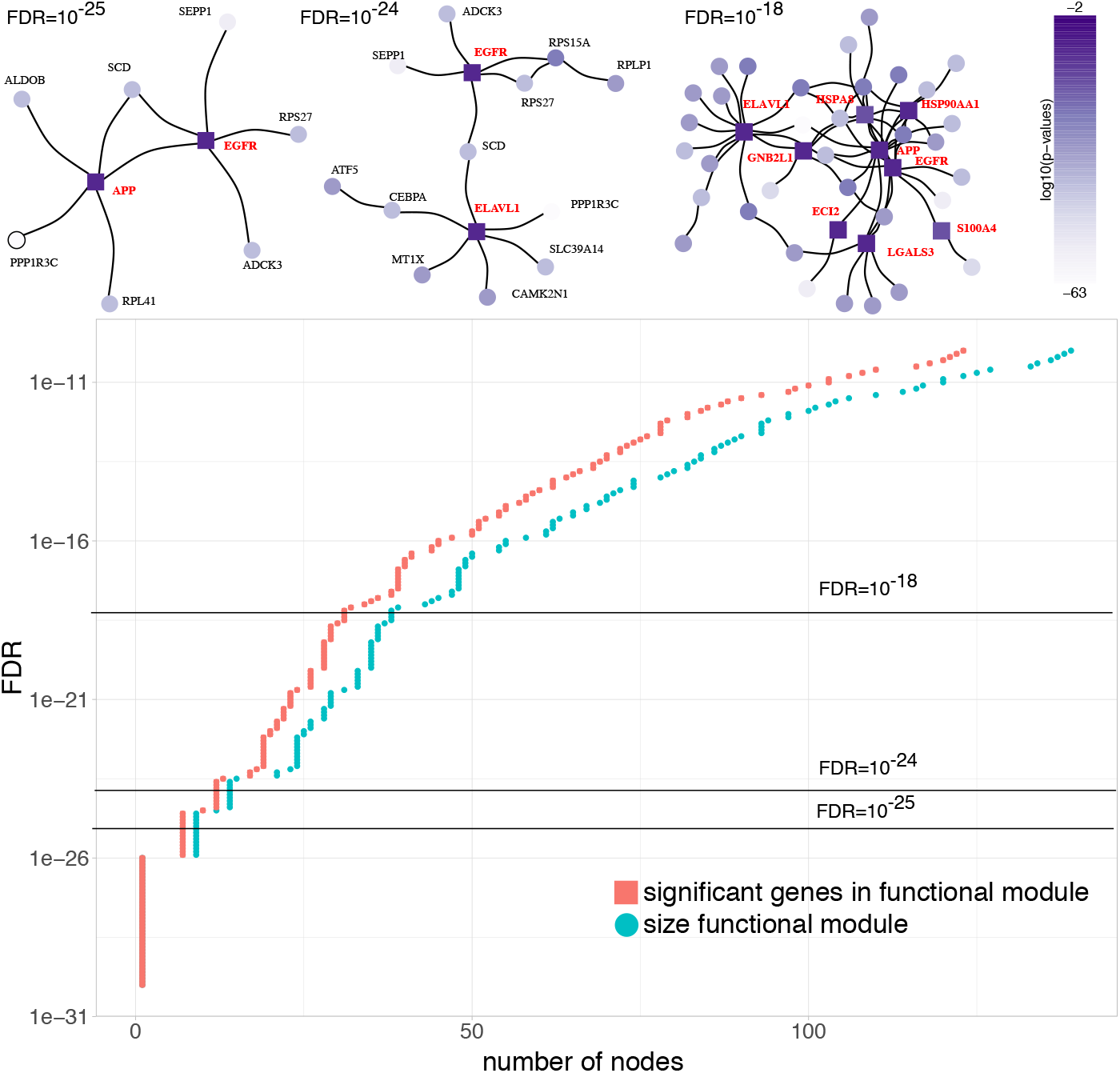
(Upper Panel) Network plots of the detected modules for three choices of the FDR (10^-25^, 10^-24^, and 10^-18^). Nodes’ colour indicates p-value from low (white) to high (purple). We indicate proteins that would not have been discovered from the gene-expression data alone as squares and give their names in bold red font. (Lower Panel) The size of detected modules (blue disks) depends on the FDR. A large fraction of proteins in these modules have significant p-values (red squares), however, for all FDR> 10^-26^, we also identify additional proteins as for a given FDR, the blue disks are to the right of the red disks.

For FDR < 10^-22^, we detect an even larger functional module, which partially overlaps with the one identified for FDR < 10^-23^, as it also includes EGFR as connector between proteins with small p-values. The second connector is *ELAV-like protein 1* (ELAVL1) with *p* ≈ 0.06. The precise function of ELAVL1 is unknown but it is believed to play a role in regulating ferroptosis in liver fibrosis [40]. For even larger FDRs, we identify a module with *M* = 42 nodes out of which 9 are not identified from the gene-expression data alone. We observe all the before-mentioned connectors, as well as, *hepatocellular carcinoma-associated Antigen 88* (ECI2) and S100 calcium-binding protein A4 (S100A4). The latter regulates liver fibrogenesis by activating hepatic stellate cells [41]. Overall, the number of proteins we identify additionally with our method is moderately increasing with the FDR. In Supplementary Note 7, we show these M(FDR) curves for all six hepotacyte clusters.

### 2.2 Functional modules for different clusters

Above, we investigated the influence of the FDR on the detected modules for a single cell cluster. As we obtained six hepatocyte clusters (see Fig. 2), we can compute DEGs and thus also active modules for each cluster separately. As each cluster represents a different cell state, different genes are identified by a DEG analysis, which then results in different functional modules. To compare the detected modules, we use scPPIN with a FDR of 10^-27^ for all of them. In Fig. 4, we show the functional modules for each of the six clusters. The detected cluster differ in size with the largest consisting of 52 nodes (cluster H2) and the smallest consisting only of a single node (Cluster H6). This occurs because the p-values of differential expression are differently distributed for each cluster. Cluster Two has the smallest p-values as its gene expression is most different from those in all other clusters, which indicates a special function of these cells in comparison to the rest. As shown for cluster H6 in Fig. 3, increasing the FDR increases also the size of the detected functional module.

**Figure 4:**
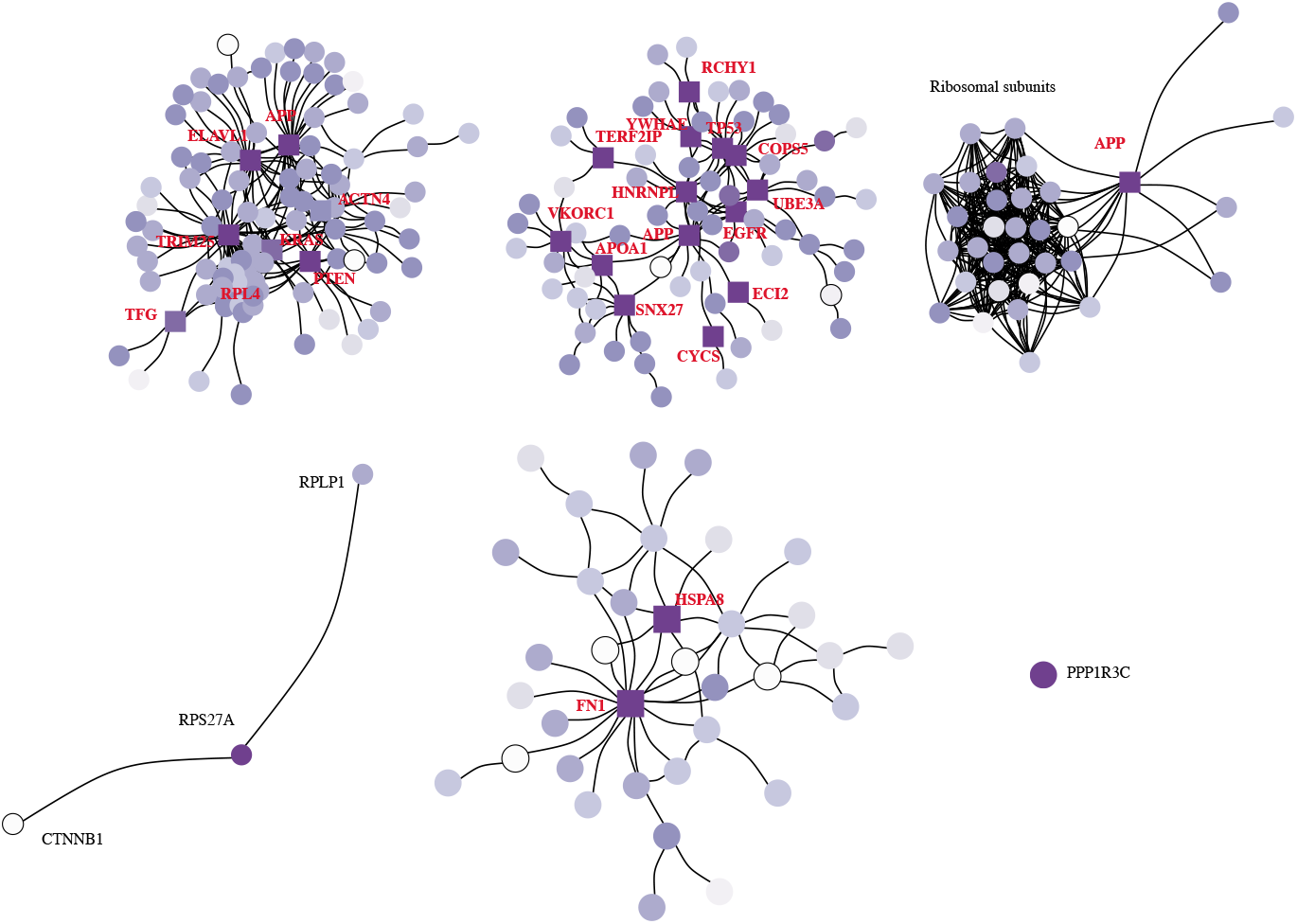
Detected modules for all six hepatocyte clusters for FDR = 10^27^. We find that the detected modules vary in size with the smallest consisting of a single protein and the largest consisting of 51 proteins. Colour indicates p-value of associated gene from low (white) to high (purple). We show nodes as squares if the could not have been detected without PPIN information. For the larger modules, we only give the names of these proteins that we would not have detected by a DEG analysis. See Supplementary Material for illustration with all protein names.

In four out of the six modules, we find proteins that we could not have identified with a DEG analysis alone. For cluster H1, these are APP, ELAVL1, TRIM25, ACTN4, PTEN, KRAS, TFG, and RPL4. For cluster H2, these are VKORC1, APOA1, SNX27, CYCS, ECI2, APP, EGFR, UBE3A, HNRNPL, COPS5, TP53, YWHAE, RCHY1, and TERF2IP. For cluster H3, this is APP. For H5, these are HSPA8 and FN1. We find that APP is identified as part of the active module in three of these clusters, which indicates that this membrane-bound protein may play an important role in different biological contexts.

To systematically access these biological contexts, we perform a GO-term enrichment test to assess the hypothesis that the detected modules represent biologically relevant pathways (see Methods). We find that all but the two smallest modules have GO terms enriched (see Table 1). The GO terms hint at distinct biological functions for the different cell clusters. Clusters H1 and H3 are involved in translational initiation, H2 in response to stress, and H5 in the extracellular structure organisation. All of these identified cellular processes represent different hepatocytes functions that have been found *in vivo* [42, 2].

**Table 1:** For each of the six clusters we give the three most enriched GO terms.

| cluster | module size $M$ | top enriched GO terms |
| --- | --- | --- |
| H1 | 51 | translational initiation<br>nuclear-transcribed mRNA catabolic process<br>SRP-dependent cotranslational protein targeting to membrane |
| H2 | 52 | response to stress<br>defense response<br>platelet degranulation |
| H3 | 27 | SRP-dependent cotranslational protein targeting to membrane<br>nuclear-transcribed mRNA catabolic process<br>translational initiation |
| H4 | 3 | *none* |
| H5 | 30 | extracellular structure organization<br>regulated exocytosis<br>alcohol metabolic process |
| H6 | 1 | *none* |

The analysis of different cell states in the scRNA-seq with scPPIN indicates that genes associated with different parts of the PPIN are active in different transcriptional states. Different biological functions of the cell clusters are reflected by different enriched GO terms. Overall, the integration of scRNA-seq with a PPIN reveals that the cells utilise the underlying PPIN differently to fulfil their diverse biological functions.

## 3 Discussion

In this study, we integrated scRNA-seq data with PPINs to construct node-weighted networks. For each cell cluster, detecting a maximum-weight connected subgraph identifies an *active module*, i.e., proteins that interact with each other and taken together the corresponding genes are significantly differently expressed. Our method scPPIN builds on advances in DEG analysis, which are standard tools for the interpretation of scRNA-seq data. As a case study, we investigated data from healthy human livers. We find that the six identified cell clusters have different subnetworks of the PPIN as functional modules in which the corresponding genes exhibiting most significantly changed expression levels. A GO-term enrichment analysis indicates that these are also associated with different biological functions. Furthermore, these subnetworks identify proteins for which the corresponding genes are not differently expressed in a given cluster but do interact with proteins for which the corresponding genes are strongly differentially expressed. These proteins are candidates for important regulatory functions in these cells. It is only through our combination of single-cell data with PPIN data that these candidate proteins can be identified. Often, they are integral membrane proteins such as FN1, EGFR, and APP, important drivers of cell fate such as P53 and KRAS, or also proteins of so-far unknown function such as TERF2IP and TFG.

In a more general setting, scPPIN can be used to systematically analyse DEG in scRNA-seq data. The identified networks that characterise each cluster help to identify and hypothesise a biological function associated to those cells. For example, we identified the gene S100A4 in the hepatocyte cluster H6, S100A4 has been identified as a key component in the activation of stellated cells in order to promote liver fibrosis [41]. Although previously identified in a population of macrophages [41, 43], we see expression of S100A4 in this cluster of hepatocytes. This indicates, that a subpopulation of hepatocytes promotes fibrogenesis in paracrine. We also identified the amyloid precursor protein (APP) and interaction partners active in multiple hepatocytes clusters. Although little is known about liver-specific functions of APP, in the central nervous system it is a key driver of Alzheimer’s disease, as source of the amyloid-β-peptide (Aβ) [44]. Due to the major role of liver in the clearance of plasma Aβ, it would be interesting to study the contribution of Aβ produced in the liver and the impact in the central nervous system. This systemic view of Alzheimer’s disease [45, 46] may reveal alternative treatments.

Despite their success, scRNA-seq techniques have methodological limitations (e.g., zero-inflation [47]). The presented technique might be further improved by considering such specific challenges, e.g., by constructing a different mixture model (see Supplementary Note 1) or implementing an imputation/noise reduction methodology.

In conclusion, we demonstrate that integrating scRNA-seq data with PPINs detects distinct enriched biological pathways and demonstrates a functional heterogeneity of cell clusters in the liver. It suggest the participation of unexpected proteins in these pathways that are undetectable from a gene-expression analysis alone. We provide an R package scPPIN, so our method can easily be integrated to current analytical workflows for single cell RNA-seq analysis.

## 4 Acknowledgements

This the work was supported by a Novo Nordisk – University of Oxford pump priming award under the Strategic Alliance between the two partners. F.K was funded through this award and is now funded by the EPSRC (funding reference EP/R513295/1). The authors would like to thank Dr. Quin F. Wills for his role that sparked this collaboration. The authors thank Lyuba V. Bozhilova for fruitful discussions.

## 5 Author contributions

E.M.T performed experiments. E.M.T, T.M., F.K. performed numerical calculations. F.K., C.M.D, and G.R. developed the statistical methods. All authors designed the study and wrote the manuscript.

## 7 Code and data availability

The scPPIN method is available as an R library under https://github.com/floklimm/scPPIN and as an online tool under https://floklimm.shinyapps.io/scPPIN-online/.

The data discussed in this publication have been deposited in NCBI’s Gene Expression Omnibus [48] and are accessible through GEO Series accession number GSE133948 (https://www.ncbi.nlm.nih.gov/geo/query/acc.cgi?acc=GSE133948).

## 8 Methods

### 8.1 Protein–Protein Interaction Network

We construct a PPIN from the publicly available BioGrid database [35], version 3.5.166. The obtained network for *Homo sapiens* has *n* = 17, 309 nodes and *m* = 296, 637 undirected, unweighted edges. While the PPIN might be directed and edge-weighted [49] (e.g., considering confidence in an interaction [50]), we consider here exclusively undirected networks without edge weights.

### 8.2 Liver Spheroid and Bioinformatics

Human primary hepatocytes from a mixture of 10 donors grown in a 3D spheroid, were purchased from InSphero AG (Switzerland) and maintained in the culture medial provided by the company. Single cell libraries were prepare with a 10X Genomics 3’ kit and sequenced in an Illumina NextSeq 500. Sequencing data demultiplexing and alignment was carried out with CellRanger with default parameters [51]. As a quality control, we only kept cells with between 500 and 6000 genes detected. A total of 2597 cells passed this quality control, of which 2123 are hepatocytes.

### 8.3 Preprocessing

We analyse the scRNA-seq data with Seurat R package v2.3.4 [52]. As a preprocessing step, we align the data with a canonical correlation analysis [52] with usage of the first nine dimensions. We identify clusters with the default resolution of one with the function *FindClusters*. To identify cell types, we use gene markers expression and in-house references datasets.

To compute a p-value of differential expression for each obtained cluster, we use the function *FindAllMarkers* with the argument Return. thresh equal to 1 and logfc.threshold set to 0.0 because we would like to obtain p-values for all genes (significant and non-significant ones). For the same reason, we do not employ a threshold for fold-change in gene expression. We exclude genes that are expressed in less than 2 % of a cluster to avoid comparing sparsely expressed genes.

### 8.4 Node-weighted network construction

The scPPIN pipeline builds on a method for the identification of functional modules as introduced by Dittrich *et al.* for analysing bulk gene-expression data [32]. Dittrich *et al.* compute maximum-weight connected subgraphs to find subnetworks that change their expression significantly in a certain disease. Here, we use a similar approach to identify subnetworks that change significantly in different clusters of cells.

Given a network *G* = {*V, E*} with node set *V* and edge set *E* ⊂ *V* × *V*, we construct a node-weighted network *G_nw_* = {*V, E, W*} by assigning each node *i* ∈ *V* a real-valued node weight *w_i_*, which we represent as a function *W*: *V* → ℝ. We construct these node-weighted networks from a PPIN and gene-expression information. The former is in the form of a network and the latter are p-values of differential expression. We assume a bijection between genes and proteins, i.e., each protein is expressed by exactly one gene, which is a simplification of the biological processes. We find this bijection by mapping GeneIDs [35].

We delete all nodes from the PPIN for which no gene-expression data is available. We present an alternative approach that can incorporate proteins with missing expression data in the Supplementary Note 4. We assign each node a score

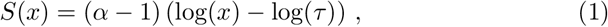

which is a function of the p-value x and we vary the *significance threshold τ* to tune the *false discovery rate* (FDR). We estimating *α* by fitting a *beta-uniform mixture model* to the observed p-values (see Supplementary Note 1). This score *S*(*x*) is negative for proteins below the significance threshold τ and positive otherwise.

### 8.5 Mathematical Optimisation Algorithm

Mathematically, the problem of identifying a subnetwork with maximal change of expression is a *maximum-weight connected subgraph problem*. Algorithmically, it is easier to solve an equivalent *prize-collecting Steiner tree* (PCST) problem [32]. Steiner trees are generalisations of spanning trees [53] and ‘prize-collecting’ indicates that the nodes have weights. To find a PCST, we use the dual ascent-based branch-and-bound framework dapcstp [54, 55]. For all calculations in this paper the algorithm identified an optimal solution in less than 10 s. For details see Supplementary Note 2.

### 8.6 Gene Ontology Enrichment

We use topGO in version 3.8 for the gene ontology enrichment (GO-enrichment) analysis. [56]. We use Fisher’s exact test to identify enriched GO terms [57]. All reported GO terms are significant with p-value 0.01 and we use a Benjamini-Hochberg procedure to counteract the multiple-comparison problem.

## Footnotes

1 If this is non-unique, multiple optimal modules of size *M* = 1 exist and can be detected. In none of our examples was this the case.

