## Supplementary Material for "Functional module detection through integration of single-cell RNA sequencing data with protein–protein interaction networks"

#### 1 Supplementary Note: Scoring Function

As in [1], we consider p-values $x \in [0, 1]$ to follow a *beta–uniform mixture* (BUM) distribution which is a mixture of noise and signal component [2]. The noise follows a uniform distribution $U([0, 1])$ with the *probability density function* (pdf)

$$f_U(x) = \begin{cases} 1\,, & \text{for } x \in [0,1] \\ 0\,, & \text{else}\,. \end{cases} \tag{1}$$

and the signal follows a beta distribution $B(\alpha, \beta = 1)$ with the pdf

$$f_B(x) = \begin{cases} \alpha x^{\alpha - 1}\,, & \text{for } x \in [0,1] \\ 0\,, & \text{else}\,. \end{cases} \tag{2}$$

The *shape parameter* $\alpha \in [0, 1]$ is a free parameter. We use $\lambda \in [0, 1]$ as *mixture parameter* to construct the distribution of p-values

$$f_{BUM}(x) = \lambda f_U(x) + (1-\lambda) f_B(x) \tag{3}$$

$$= \begin{cases} \lambda + (1-\lambda)\alpha x^{\alpha-1}\,, \ \text{for } x \in [0,1] \\ 0\,, \ \text{else}\,. \end{cases} \tag{4}$$

For each run of our algorithm, we obtain estimates of shape parameter $\alpha$ and mixture parameter $\lambda$ by performing a maximum-likelihood estimation. For this, we use an iterative limited-memory Broyden-Fletcher-Goldfarb-Shanno algorithm [3].

Fig. 1 shows a histogram of the p-values for the one-vs-one comparison between clusters H1 and H3 and the fitted distribution. Fitting a BUM model yields a mixture parameter of $\lambda \approx 0.549$ and a shape parameter of $\alpha \approx 0.482$. The BUM is in decent agreement with the observed p-values (a one-sample Kolmogorov–Smirnov test yields a statistic of $D \approx 0.014$ and a p-value of 0.086). We calculate the value of the pdf at $x = 1$, which represents the background noise, as

$$\tilde{f} = f_{BUM}(x = 1) = \lambda + (1-\lambda)\alpha \approx 0.77\,, \tag{5}$$

and show is a dotted horizontal line in Fig. 1.

We use an approach as outlined in [1] to construct an additive score $S(x) \in \mathbb{R}$ with negative values indicating background noise and positive values representing signal. This allows us to compute the differential expression of a set of genes by adding their associated scores. An appropriate choice is

$$S(x) = (\alpha - 1)\left(\log(x) - \log(\tau)\right)\,, \tag{6}$$

where we consider p-values below the *significance threshold* $\tau > 0$ as noise and those above as signal. As shown in [2], the significance threshold is a function of the FDR and we may estimate it for a BUM as

$$\tau(\text{FDR}) = \left(\frac{\tilde{f} - \text{FDR}\cdot\lambda}{\text{FDR}(1-\lambda)}\right)^{1/(\text{FDR}-1)} \quad \text{with } \tilde{f} = f_{BUM}(x=1) = \lambda + (1-\lambda)\alpha\,. \tag{7}$$

To summarise, the procedure for computation of the node scores from p-values is as follows:

1. estimation of mixture parameter $\lambda$ and shape parameter $\alpha$ from the distribution of p-values,
2. choice of an FDR as appropriate for data of choice,
3. computation of the the appropriate significance threshold through Eqn. (7), and
4. computation of the node scores through Eqn. (6).

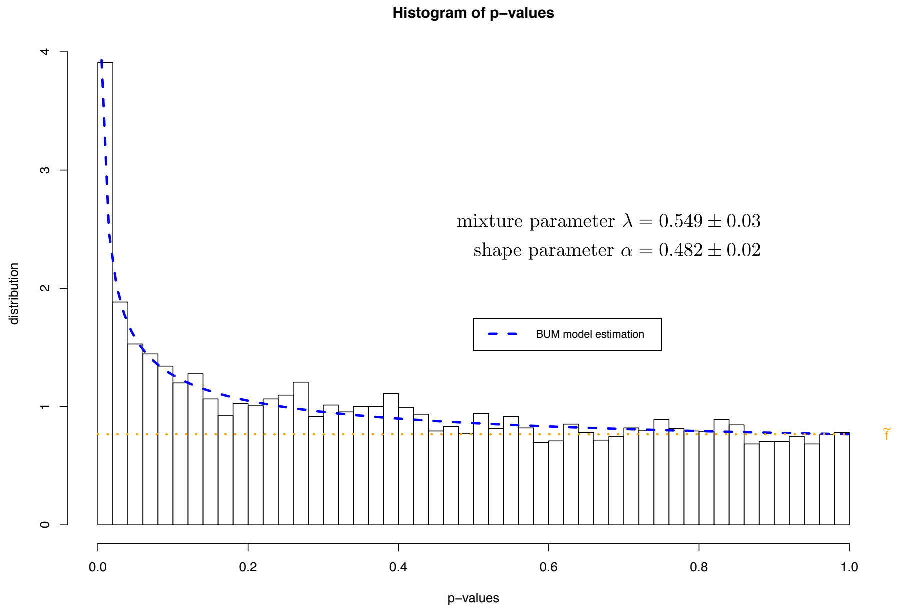


Figure 1: Fit of the BUM model to p-values for comparison cluster H1 vs cluster H3. We show a histogram of the observed p-values and the fitted BUM model (blue dashed curve). The mixture parameter is $\lambda \approx 0.549$ and the shape parameter of the beta distribution is $\alpha \approx 0.482$. We show the probability $\tilde{f} = f(x = 1) \approx 0.77$ as a horizontal dotted orange line.

### 2 Supplementary Note: Optimisation

As outlined in Supplementary Note 1, we assign each gene a score $S \in \mathbb{R}$. Under the null hypothesis, these scores are independent and identically distributed random variables. For any set $Q$ of genes, we can compute the score

$$S_Q = \sum_{i \in Q} S(x_i) \,, \tag{8}$$

which indicates to what extent this set $Q$ is significantly differently expressed.

The aim of our optimisation algorithm is the identification of a set of genes that maximises this score (i.e., being strongly differently expressed in a module) while the associated proteins are connected to each other in the PPIN. Mathematically, this problem is known as a *maximum-weight connected subgraph* (MWCS) problem.

**Problem.** (Maximum-weight connected subgraph, MWCS) *Given a vertex-weighted graph $G = (V, E, W)$, find a connected subgraph $T = (V_T, E_T, W) \subset G$ that maximises the score $W(T) = \sum_{i \in V_T} W(i)$.*

Here, we use $T \subset G$ to indicate that $T = (V_T, E_T)$ is a subgraph of $G$, i.e., $V_T \subset V$ and $E_T \subset E$. A graph is connected if there is a path between every pair of edges [4].

Finding a MWCS is NP-hard. A heuristic approach was used in [5] to identify the MWCS. It is computationally advantageous to transform this problem into an equivalent *prize-collecting Steiner tree* (PCST) problem because it is often possible to obtain provably optimal solutions for this transformed instance [1, 6].

**Problem.** (Prize-collecting Steiner tree, PCST) *Given a graph* $G = (V, E, c, p)$ *in which the edge weights* $c : E \to \mathbb{R}^{\geq 0}$ *indicate costs and the node weights* $p : V \to \mathbb{R}^{\geq 0}$ *indicate profits, find a connected subgraph* $T \subset G$ *that maximises the profit*

$$p(T) = \sum_{i \in V_T} p(v) - \sum_{e \in E_T} c(e)\,.$$

We can identify a MWCS by transforming the node-weighted network $G = (V, E, W)$ into a network $G = (V, E, c, p)$ with non-negative node and edge weights. Specifically, we compute $p(i) = w(i) - w_{\min}$ and $c(e) = -w_{\min}$, where $w_{\min} = \min_{i \in V(G)} w_i$, so that all edges have the same weight. See Fig. 2, for an example and [1] for a proof of this equivalence.

Here, we use the dual ascent-based branch-and-bound framework DAPC-STP [7, 8] to find the PCST. While this algorithm is not guaranteed to find an optimal solution, in practice it finds solutions close to optimality for networks with hundreds of thousands of nodes and millions of edges in minutes to hours. For all computations on the PPINs in this manuscript, we found optimal solutions in less than ten seconds.

PCSTs are always trees (i.e., if they have $N$ nodes they have $N - 1$ edges) because adding additional edges to the PCST decreases the profit $P$. Accordingly, MWCSs are also trees. When visualising the active modules, however, we construct a *node-induced subgraph*, which is the set of nodes in $T$ with all edges from $G$. Therefore the modules may contain loops.

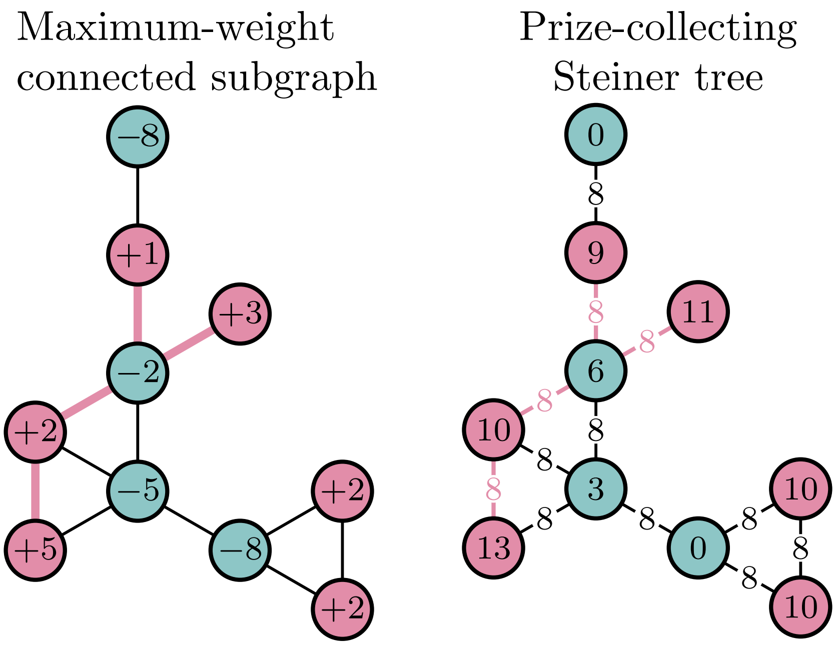


Figure 2: Example of a node-weighted network for which we want to find the MWCS (left) and its equivalent node- and edge-weighted network for which we find the PCST (right). We transform this problem by shifting all node weights by $w_{\min} = -8$ and assigning each edge a cost of $c = -w_{\min} = 8$. We show nodes with a positive score in the node-weighted network in red and nodes with negative score in blue. We highlight the optimal solution in both instances by colouring the edges red. The MWCS has a weight of $W = 5{+}2{-}2{+}3{+}1 = 9$ and the PCST has a profit of $P = 13+10+6+11+9-4\times 8 = 49-32 = 17 = 9+8$.

#### 2.1 Pairwise comparison between cell clusters

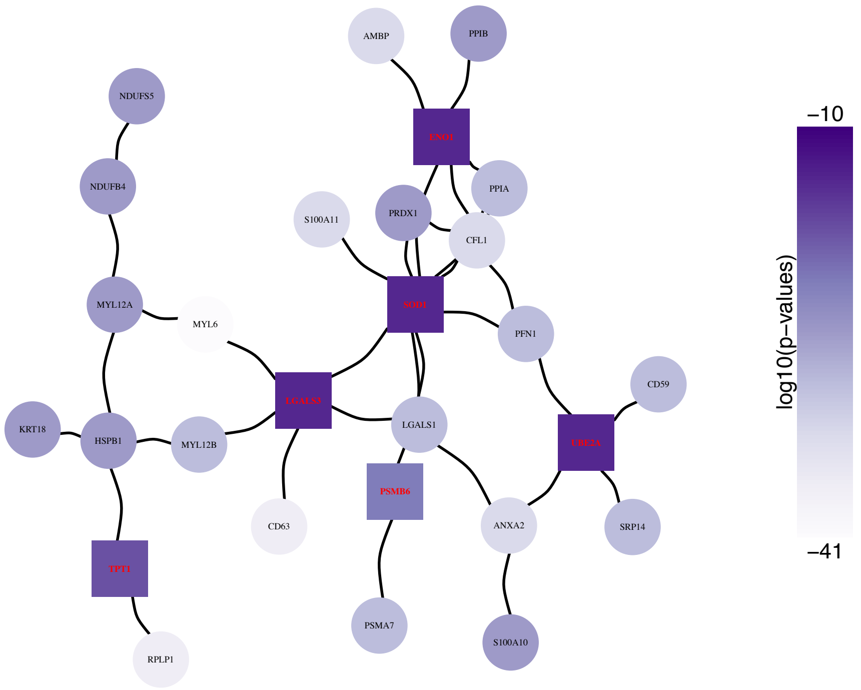


Figure 3: We can compare also the expression between two clusters. Here, we show the active module in cluster H2 vs cluster H4.

In the main manuscript, we have identified active modules in cell clusters by comparing the expression of genes against the expression of all other cells (‘one-vs-all’). It is also possible to compare the expression of any two clusters pairwise (’one-vs-one’) and obtain a potentially more nuanced picture of two clusters that have a similar but distinct transcriptional state.

In Fig. 3, we show the detected functional module in a pairwise comparison of clusters H2 and H4. To compute the p-values of differential expression between two clusters, we can also use the FindMarkers function in Seurat. The rest of the analysis pipeline in scPPIN is the same. We identify a module that consists of proteins such as TPT1, a tumour protein, and SOD1, which is involved in apoptosis.

We compare cluster H1 vs H3 and detect a functional module as shown in Fig. 4. Note that the FDR$=0.01$ is very large (in comparison with earlier calculations and FDR$=10^{-19}$ in the main manuscript) as both clusters have a very similar gene expression. A GO-term enrichment analysis finds only the term ’regulation of symbiosis, encompassing mutualism through parasitism’ enriched. Our method identifies TRIM25, an E3 ubiquitin ligase enzyme [9]. TRIM25

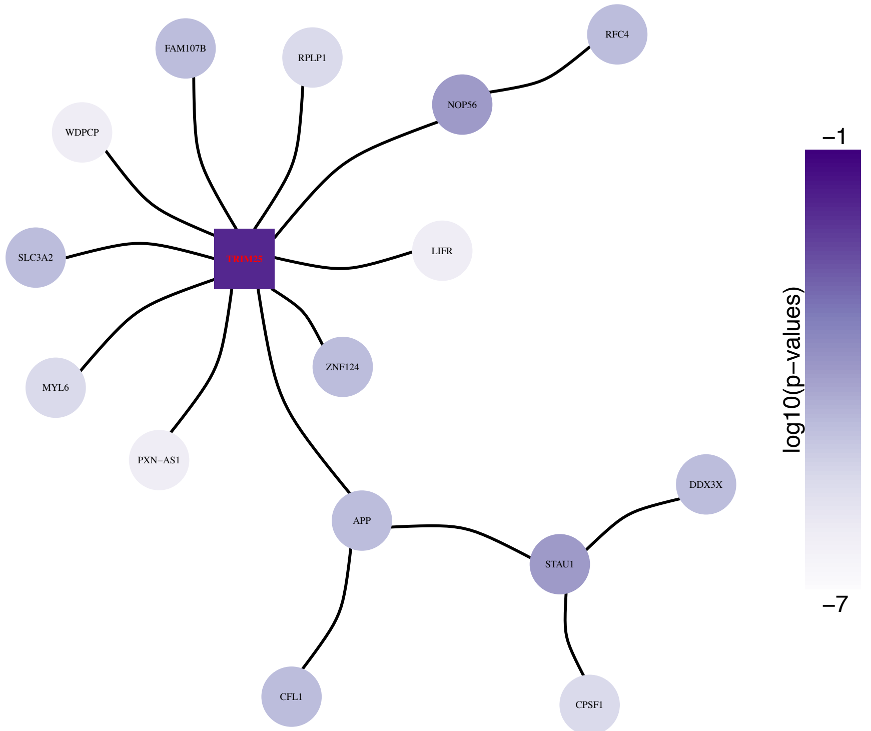


Figure 4: We can compare also the expression between two clusters. Here, we show the active module in cluster H1 vs cluster H3.

binds to viral RNA and so regulates immune response against viruses. These results indicate that either: one of the clusters is infected with a virus or this clusters shows a response to inflammation that is using pathways that are usually associated with antiviral response.

### 3 Supplementary Note: Missing expression data

In the method described in the main manuscript and Supplementary Note 1, we assign each node in a PPIN a score based on the p-value of differential expression. This is only possible if we have the gene-expression measurement for a given protein. In practice, we delete nodes that represent proteins without available expression value. One alternative approach is to assign each node in a PPIN for which we have no expression information of the corresponding gene, a small negative score (i.e., $S = S_0 = -1$). There is no *a priori* correct way to choose $S_0$ and smaller value decrease the likelihood that such proteins are part of the detected functional module. For $S_0 \to -\infty$, we recover the case without missing-data replacement.

In Fig. 5, we show a detected functional module for a node-weighted PPIN in which we assigned proteins without gene expression data $S_0 = -1$. It is the same underlying data and FDR = 0.01 as in Fig. 4. As before, we detect TRIM25 and interacting proteins. In addition, we also detect three proteins without expression information, RPA2, RBM4, and SOX2. The total number of proteins in this active module increases from 16 to 20. Note that for some of these additional proteins (e.g., JUND and RPP40), we have gene expression data but they were not in the MWST. We now detect them as part of the functional module because they are connected to the original module via the missing-data nodes.

This example demonstrates that incorporating nodes with no associated scRNA-seq data may alter the detected modules. The choice of $S_0$ is a free parameter and a principled approach for detecting them will be part of future research.

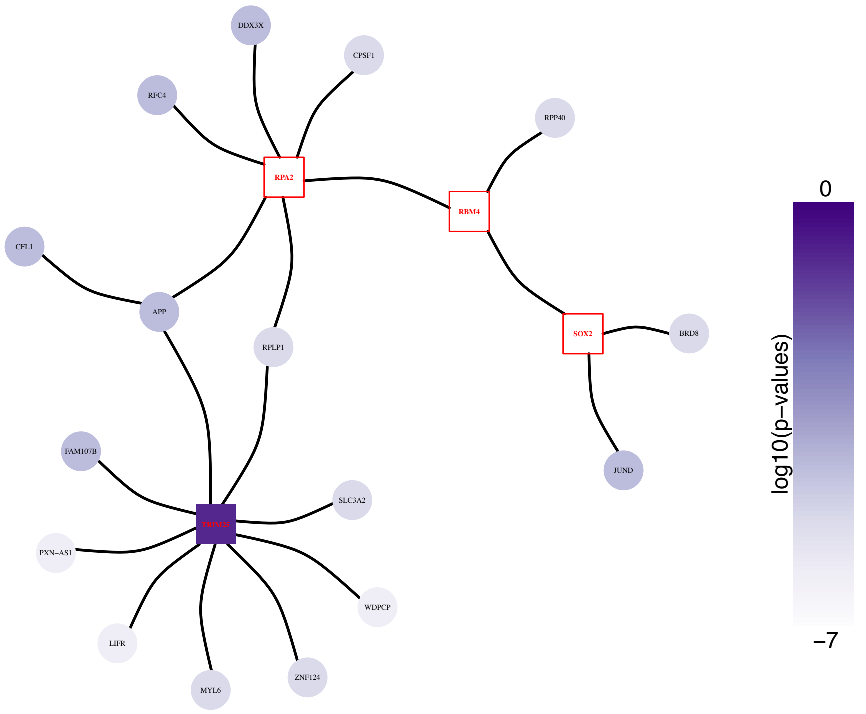


Figure 5: Assigning proteins without expression data for corresponding genes a score $S_0$ allows to keep these proteins in the node-weighted PPIN. Here, we show the active module in cluster H1 vs cluster H3 for $S_0 = -1$. In comparison with Fig. 4, we detect here three proteins without expression information: RPA2, RBM4, and SOX2 (shown as red boxes).

### 4 Supplementary Figure: Detected modules for all six hepatocyte clusters

Fig. 6 shows the detected modules for all six hepatocyte clusters FDR $= 10^{27}$. It is identical to Fig. 4 in the main manuscript but with all proteins labelled.

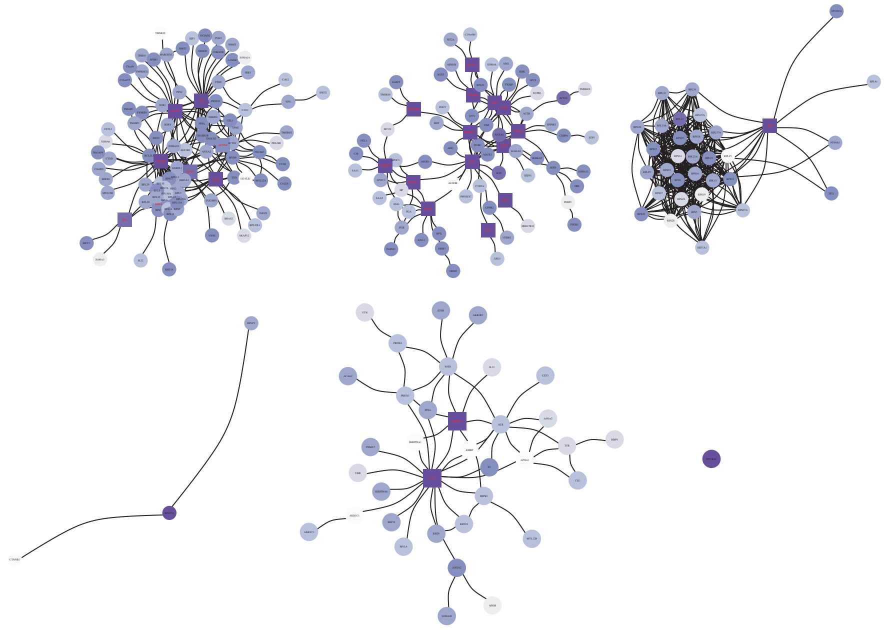

Figure 6: Detected modules for all six hepatocyte clusters for FDR $= 10^{27}$. We find that the detected modules vary strongly in size with the smallest consisting of a single protein and the largest consisting of 51 proteins. Colour indicates p-value of associated gene from low (white) to high (purple). We show nodes as squares if the could not have been detected without PPIN information.

### 5 Supplementary Note: Influence of FDR on size of detected modules

In the main manuscript, we show the influence of the FDR on the size of the detected module for one cluster. Here, we show this for all six clusters. For all clusters we compute the functional modules for FDR $\in [10^{-45}, 10^{-15}]$. Note that the horizontal axes are scaled differently for each cluster for illustration purposes.

For all clusters, we find that the size of the functional modules is increasing with the FDR. Furthermore, for all clusters, there are FDR-choices for which we detect modules that have some proteins that we would not have detected from a DEG analysis alone.

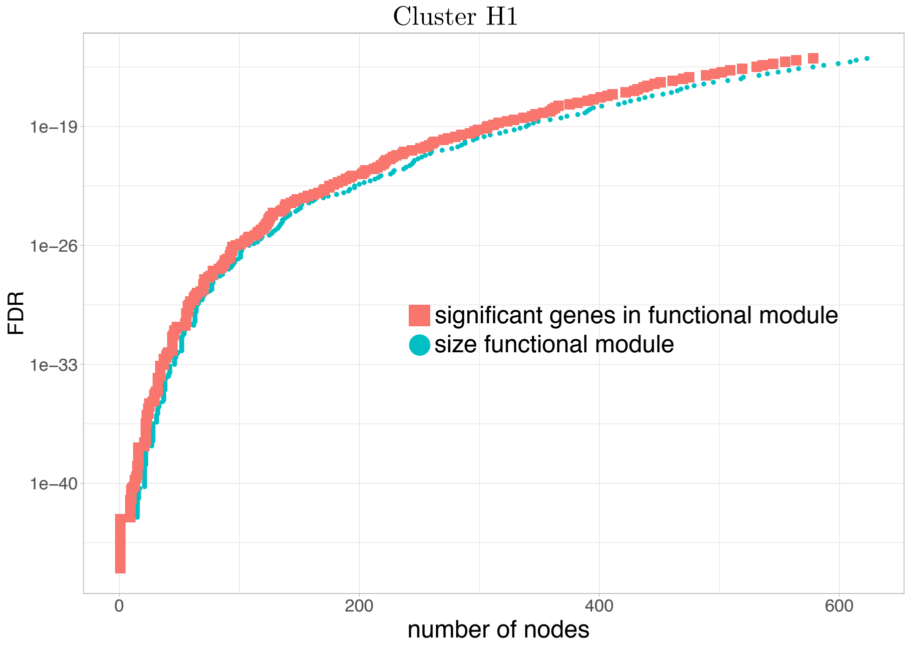


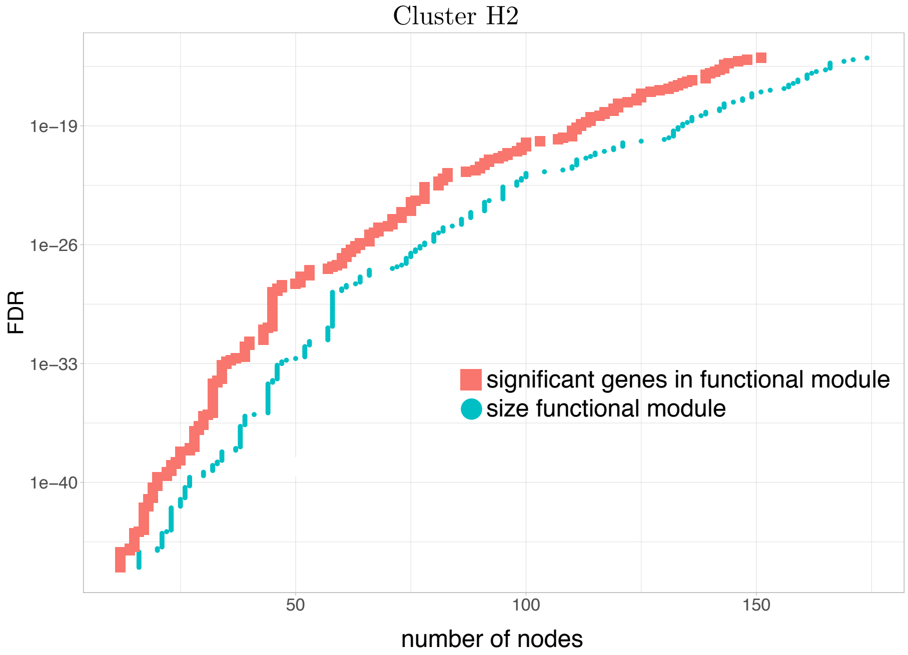


Figure 7: The module size as a function of the FDR for clusters H1 and H2.

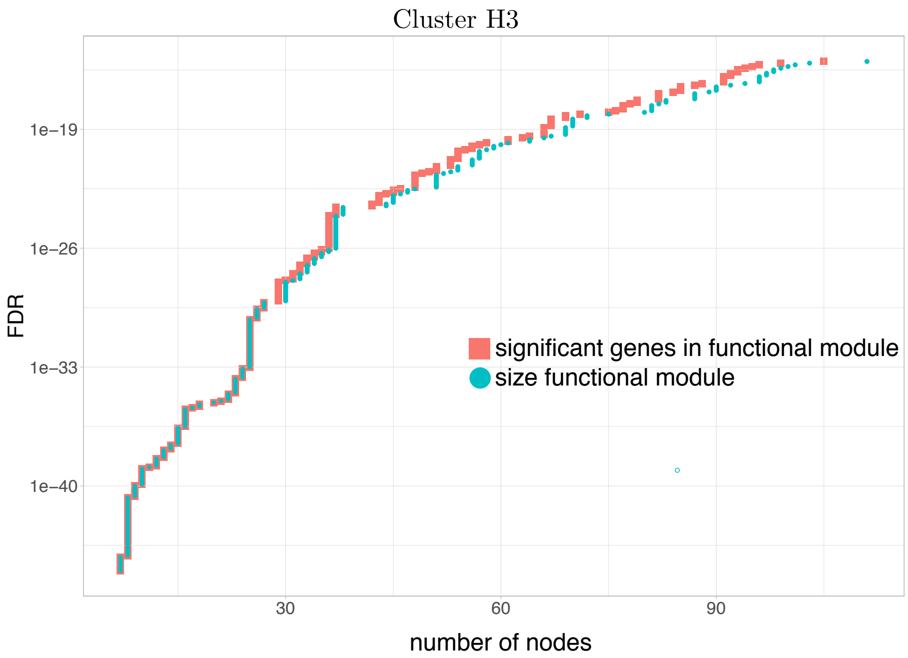


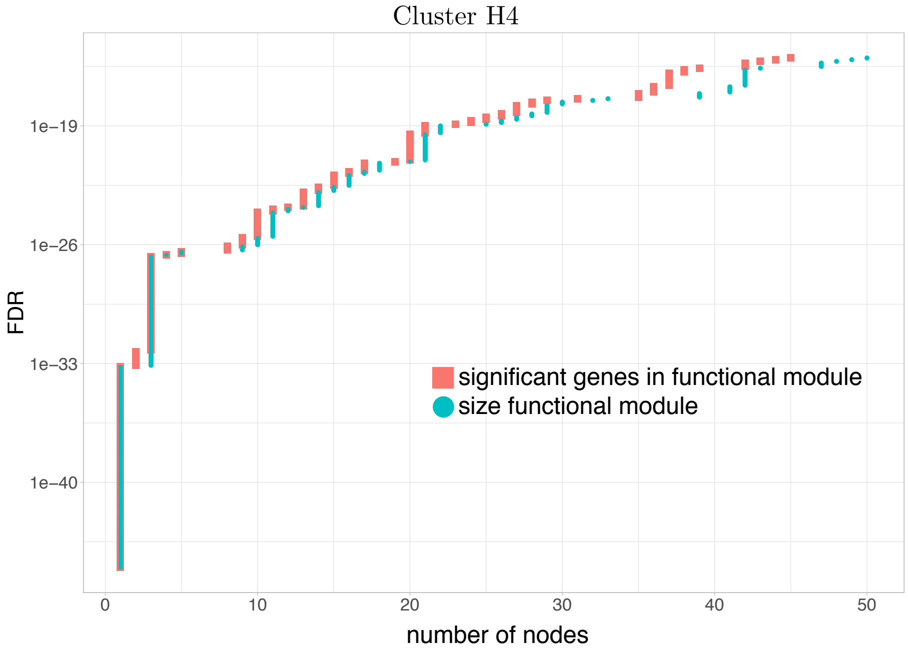


Figure 8: The module size as a function of the FDR for clusters H3 and H4.

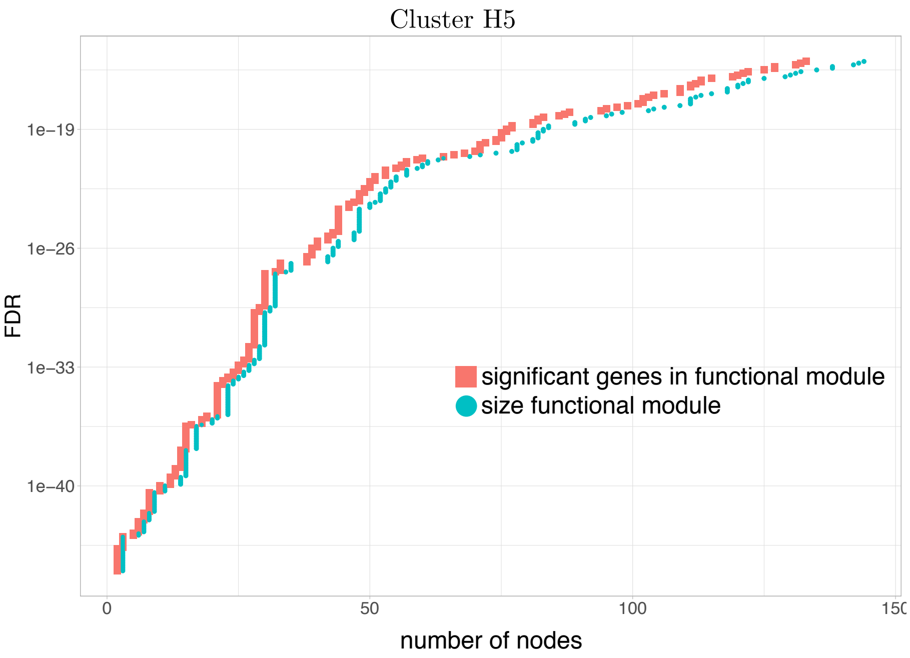


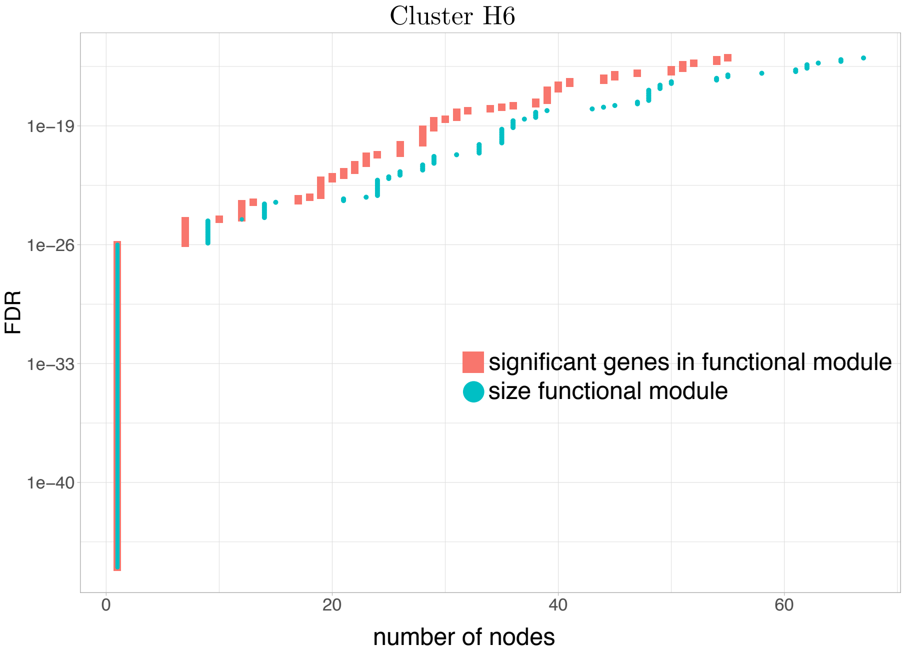


Figure 9: The module size as a function of the FDR for clusters H5 and H6.

[4] Geir Agnarsson and Raymond Greenlaw. *Graph Theory: Modeling, Applications, and Algorithms*. Prentice-Hall, Inc., 2006.

[5] Trey Ideker, Owen Ozier, Benno Schwikowski, and Andrew F Siegel. Discovering regulatory and signalling circuits in molecular interaction networks. *Bioinformatics*, 18(suppl_1):S233–S240, 2002.

[6] Ivana Ljubić, René Weiskircher, Ulrich Pferschy, Gunnar W Klau, Petra Mutzel, and Matteo Fischetti. An algorithmic framework for the exact solution of the prize-collecting steiner tree problem. *Mathematical Programming*, 105(2-3):427–449, 2006.

[7] Matteo Fischetti, Markus Leitner, Ivana Ljubić, Martin Luipersbeck, Michele Monaci, Max Resch, Domenico Salvagnin, and Markus Sinnl. Thinning out steiner trees: a node-based model for uniform edge costs. *Mathematical Programming Computation*, 9(2):203–229, 2017.

[8] Markus Leitner, Ivana Ljubić, Martin Luipersbeck, and Markus Sinnl. A dual ascent-based branch-and-bound framework for the prize-collecting steiner tree and related problems. *INFORMS Journal on Computing*, 30(2):402–420, 2018.

[9] María Martín-Vicente, Luz M Medrano, Salvador Resino, Adolfo García-Sastre, and Isidoro Martínez. TRIM25 in the regulation of the antiviral innate immunity. *Frontiers in Immunology*, 8:1187, 2017.
